# Circulating CD138 (syndecan-1) enhances APRIL-mediated autoreactive B cell survival and differentiation in MRL/Lpr mice

**DOI:** 10.1101/2021.05.11.443667

**Authors:** Lunhua Liu, Mustafa Akkoyunlu

## Abstract

High levels of serum CD138, a heparan sulfate-bearing proteoglycan, correlates with increased disease activity in systemic lupus erythematosus (SLE) patients. Mechanisms responsible for serum CD138 production and its biological function in SLE disease remain poorly understood. In this study, corroborating patient data, we detected an increase in serum CD138 in MRL/Lpr mice parallel to disease activity. Although TCRβ+CD138+ T cells expand in MRL/Lpr mice as the disease progresses, surprisingly, TCRβ+CD138- cells were the primary source of circulating CD138 as the transfer of TCRβ+CD138- cells to young MRL/Lpr mice, but not TCRβ+CD138+ cells, resulted with higher serum CD138 in the recipient mice. We found that elevated trypsin, expressed by TCRβ+CD138- cells, was able to cleave CD138 from T cells. Moreover, suggesting the contribution of cleaved CD138 to the increase in blood CD138, trypsin inhibitors ‘defined trypsin inhibitor’ (DTI) or leupeptin increased CD138 expression on TCRβ+CD138- cells. Furthermore, soluble CD138 was able to bind ‘a proliferation inducing ligand’ (APRIL) and enhanced APRIL-mediated plasma cell generation and autoreactive antibody production through the phosphorylation of extracellular-signal-regulated kinase (ERK) in B cells. APRIL receptor, ‘transmembrane activator, calcium modulator, and cyclophilin ligand interactor’ (TACI) was involved in the enhancement of APRIL activity by CD138, as the synergistic effect of APRIL and CD138 was ablated on TACI deficient B cells. These findings indicate a regulatory role for soluble CD138 in B cell differentiation and autoreactive antibody secretion in SLE disease.

## Introduction

Systemic lupus erythematosus (SLE) is a chronic autoimmune disease characterized by hyperproduction of autoreactive antibodies that cause inflammation and multiple organ damage (1). This systemic pathological immune response, which involves both innate and adaptive immune system, is characterized by the elevation of multiple cytokines in serum. Increased serum levels of ‘A Proliferation-Inducing Ligand’ (APRIL), ‘B-cell-activating factor’ (BAFF), IFN-α, IFN-γ, IL-6, IL-12, IL-17 and TNF-α are found to be positively correlated with autoreactive antibody production, SLE Disease Activity Index (SLEDAI) scores, and organ involvement (2). Among these cytokines, APRIL and BAFF bind to B cell maturation antigen (BCMA) and transmembrane activator, calcium modulator, cyclophilin ligand interactor (TACI) and BAFF additionally binds to BAFF receptor (BAFFR) (3). Engagement of these three receptors can directly lead to the maturation, proliferation and survival of B cells in addition to the formation of long lived antibody secreting plasma cells (3). Thus, considerable interest has been expressed in the development of APRIL and BAFF antagonists as therapeutic agents. A fully humanized anti-BAFF monoclonal antibody (belimumab) which effectively reduces disease activity and flare severity has been approved in the United States, and around the world for the treatment of lupus (4).

In addition to the increase in inflammatory cytokines, SLE patients, but not rheumatoid arthritis patients, manifest with elevated serum levels of CD138 (syndecan-1) (5,6). As a member of the syndecan family of type I transmembrane proteoglycans, CD138 is composed of a core protein modified by heparan sulphate and chondroitin sulphate chains (7,8). Membrane bound CD138 has been shown to play an important role in wound healing, cell adhesion, and endocytosis (9). Its biological activity is mediated by the binding of its covalently attached glycosaminoglycan chains to several extracellular adhesion molecules, growth factors, cytokines and chemokines (7,10). The attachment of CD138 to these molecules results in the modification of their biological activity (7,10).

Like the other three members of syndecan family molecules, the intact ectodomain of CD138 is constitutively shed from cells and forms soluble CD138 as part of normal cell surface heparan sulfate proteoglycan turnover (11,12). In response to injury or infection, the ectodomain of CD138 is proteolytically shed from the cell surface by matrix metalloproteinases (MMP), such as MMP9 and matrilysin (MMP7) (11,13). Elevated soluble CD138 also regulates a variety of pathways that are related to wound healing, cell proliferation and apoptosis (14). Excessive soluble CD138 has been shown to delay skin wound repair as it enhances elastase activity and inhibits growth factor action (15). Soluble CD138 is also involved in host response to tumor development. For example, soluble CD138 inhibits the mitogenicity of fibroblast growth factor 2 (FGF-2) on B stem-like F32 cells and overexpression of soluble ectodomain or the addition of exogenous CD138 ectodomain significantly inhibits the proliferation of MCF-7 breast cancer cells line (16,17). Moreover, the impact of shed CD138 extends beyond cell to cell contact within the tumor microenvironment as the accumulation of soluble CD138 enhances the growth of myeloma tumors in vivo and promotes endothelial invasion and angiogenesis (18–20). Besides, in myeloma and lung cancer patients, high levels of serum CD138 correlates with poor disease prognosis and survival (21,22).

In SLE patients, serum CD138 levels positively correlate with SLEDAI and anti-dsDNA antibody levels (5,6). But the origin and function of circulating CD138 in lupus patients remain largely unknown. In this study, we investigated the origin and biological function of soluble CD138 in lupus development. We first focused on TCRβ+CD138+ cells as the source of soluble CD138 because we have recently reported the expansion of CD138 bearing TCRβ+ cells in various organs of the lupus prone MRL/Lpr mouse (23). Surprisingly, we found that activated TCRβ+CD138- cells produce more soluble CD138 than activated TCRβ+CD138+ cells. Moreover, the transfer of TCRβ+CD138- cells into MRL/Lpr mice led to higher serum CD138 measurement then the transfer of TCRβ+CD138+cells did. In support of TCRβ+CD138- cells as the source of circulating CD138, we found higher expression of trypsin by TCRβ+CD138- cells then TCRβ+CD138+ cells, which effectively cleaved CD138 to produce its soluble form. Interestingly, we also found that binding of soluble CD138 to APRIL strongly enhanced APRIL induced extracellular-signal-regulated kinase (ERK) phosphorylation in B cells and promoted B cell differentiation into antibody secreting plasma cells.

## Results

### Activated TCRβ+CD138- cells release more soluble CD138 than TCRβ+CD138+ cells do

SLE patients manifest with increased serum CD138 levels, which correlate with disease activity and severity of nephritis (5,6). By using the widely studied lupus prone MRL/Lpr mice, we investigated the source of CD138 in lupus disease (24). In MRL/Lpr mice, a single mutation in the *fas* apoptosis gene results in lymphoproliferation and autoreactive B and T cell activation (25). As a result, MRL/Lpr mice begin to manifest lupus symptoms such as anti-dsDNA antibodies and kidney disfunction starting from 4 to 6 weeks of age, and the diseases progresses with age (25,26). Analysis of serum CD138 levels in MRL/Lpr mice from different ages indicated that the serum CD138 levels increase with age (Fig. 1*A*). Thus, as reported in lupus patients, serum CD138 levels positively correlate with the disease progression in MRL/Lpr mice. We have recently reported the expansion of TCRβ+CD138+ cells in MRL/Lpr mice (23). Spleens of 10 weeks old MRL/Lpr mice harbor CD138-expressing plasmablasts, plasma cells and TCRβ+CD138+cells (Fig. 1*B*, Fig. S1). As the disease progresses, both the CD138-expressing plasma cells and TCRβ+CD138+ T cells expand in the blood also (Fig. 1*C*). Importantly, the numbers and percentages of TCRβ+CD138+ were significantly higher than the plasma cells in both the spleen and blood (Fig. 1*B*, *C*). Thus, we speculated that the shedding of CD138 from TCRβ+CD138+ cells may be responsible for the elevated circulating CD138. To test this hypothesis, we adoptively transferred TCRβ+CD138+ or TCRβ+CD138- cells from 12 to 14 weeks old MRL/Lpr mice into 8 weeks old MRL/Lpr mice and measured serum CD138 levels in the recipient mice. Surprisingly, analysis of serum CD138 levels 3 days after the transfer indicated that the TCRβ+CD138- cell-transferred group had higher serum CD138 levels than PBS and TCRβ+CD138+ cell-transferred groups (Fig. 1*D*). Corroborating the adoptive transfer experiment results, when stimulated with anti-CD3 and - CD28 antibodies (αCD3/CD28), purified TCRβ+CD138- secreted more CD138 than activated TCRβ+CD138+ cells did (Fig. 1*E*). These data suggested that TCRβ+CD138- cells may be an important source of circulating CD138 in MRL/Lpr mice.

**Figure 1.**
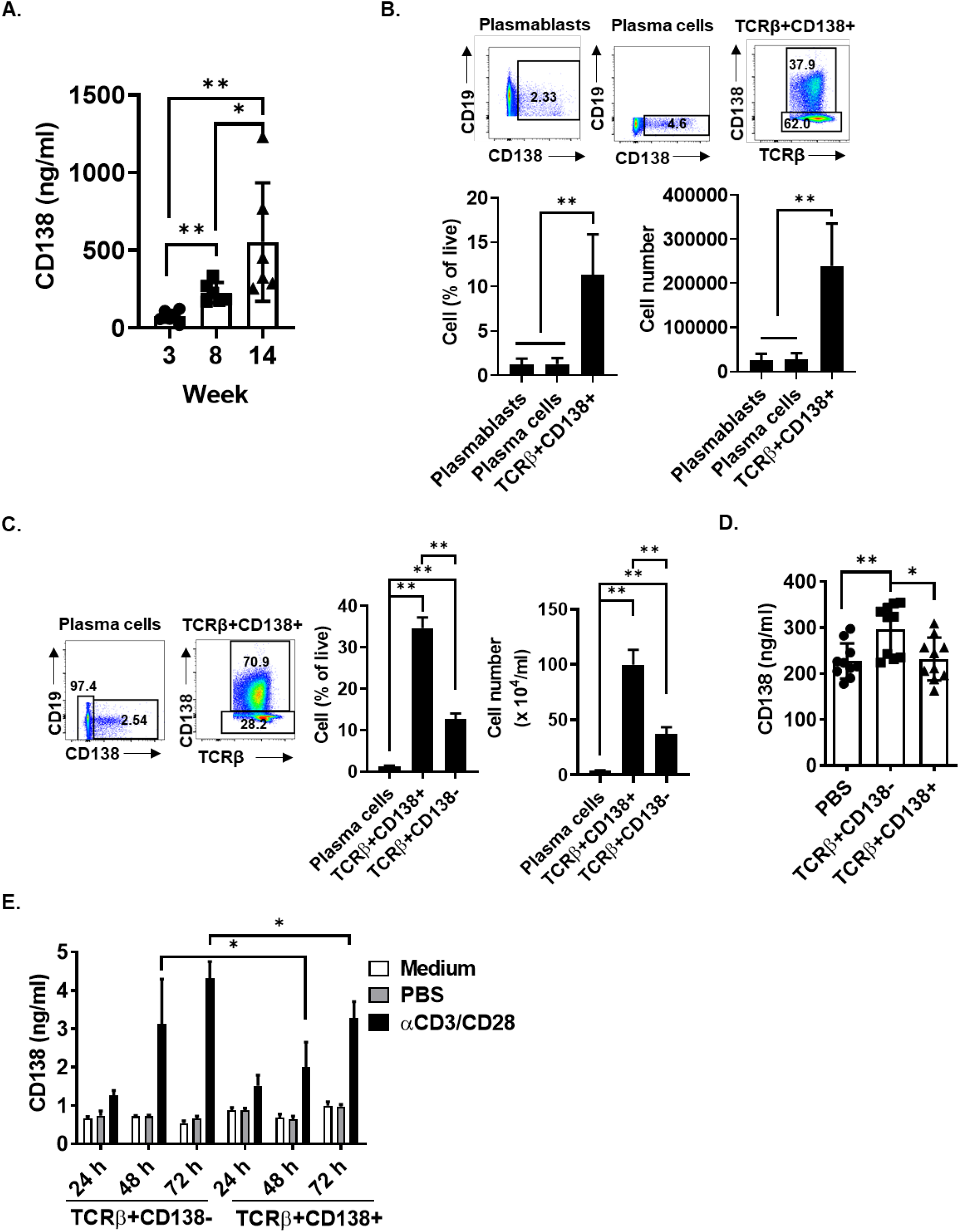
Activated TCRβ+CD138- cells secrete more CD138 than TCRβ+CD138+ cells do. ***A***, serum CD138 levels in MRL/Lpr mice from different ages were measured by ELISA. Mean ± SD of 6 mice from two independent experiments are plotted. ***B***, splenic plasmablasts (CD19+CD138+), plasma cells (CD19-CD138+) and TCRβ+CD138+ cells were characterized and enumerated from 10 weeks old MRL/Lpr mice. Mean ± SD of 5 mice from two independent experiments are plotted. ***C***, blood was collected from 14 weeks old MRL/Lpr mice, and plasma cells (CD19-CD138+), TCRβ+CD138+ cells, and TCRβ+CD138- cells were quantified in FACS. Mean ± SD of 5 mice from two independent experiments are plotted. ***D***, purified TCRβ+CD138- or TCRβ+CD138+ cells from 12 to 14 weeks old MRL/Lpr mice were adoptively transferred into 8 weeks old MRL/Lpr mice and the serum CD138 levels were determined by ELISA 3 days after the transfer. Mean ± SD of 10 mice from two independent experiments are plotted. ***E***, purified TCRβ+CD138- and TCRβ+CD138+ cells were activated for indicated duration and culture supernatant CD138 levels were measured by ELISA. Mean ± SD of three independent experiments are plotted. *p<0.05, **p<0.01.

### CD138 is cleaved from lupus T cells by trypsin

Next, we sought to determine how membrane bound CD138 was released from lupus T cells. In a variety of cells ranging from epithelial cells, macrophages and CD4+ T cells to cells from tumors such as myeloma, pancreatic cancer and melanoma, membrane CD138 is cleaved by different proteinases such as MMP9, MT1-MMP, MT3-MMP, collagenase, and trypsin (11,13,19,27–30). We therefore tested the activity of these proteinases on the cleavage of CD138 from lupus mouse T cells. We found that MMP9 was unable to cleave CD138 from TCRβ+CD138+ cells (Fig. 2*A* and Fig. S2*A*, *B*). Similarly, collagenase I and D failed to shed CD138 from purified TCRβ+CD138+ cells (Fig. 2*B*). In contrast to MMP9 and collagenases I and D, we found that CD138 was highly sensitive to trypsin cleavage as after a short term (5-minute) treatment with trypsin or TrypLe, cell surface CD138 expression sharply decreased on TCRβ+CD138+cells (Fig. 2*C*, and Fig. S2*C*, *D*). Supporting the observed trypsin activity, analysis of mouse CD138 amino acid sequence indicated two trypsin targets, arginine and lysine amino acids located at the C-terminal of CD138 extracellular domain (Fig. S2*E*). To verify the trypsin cleavage and release of CD138 from TCRβ+CD138+ cells, we measured the CD138 content in the supernatants from short-term trypsin treated cells by ELISA. Compared to the supernatants from untreated cells, significantly higher levels of soluble CD138 was detected in the supernatants of trypsin treated cells (Fig. 2*D*). Thus, CD138 on lupus T cells is highly sensitive to trypsin cleavage but resistant to MMP9 and collagenases.

**Figure 2.**
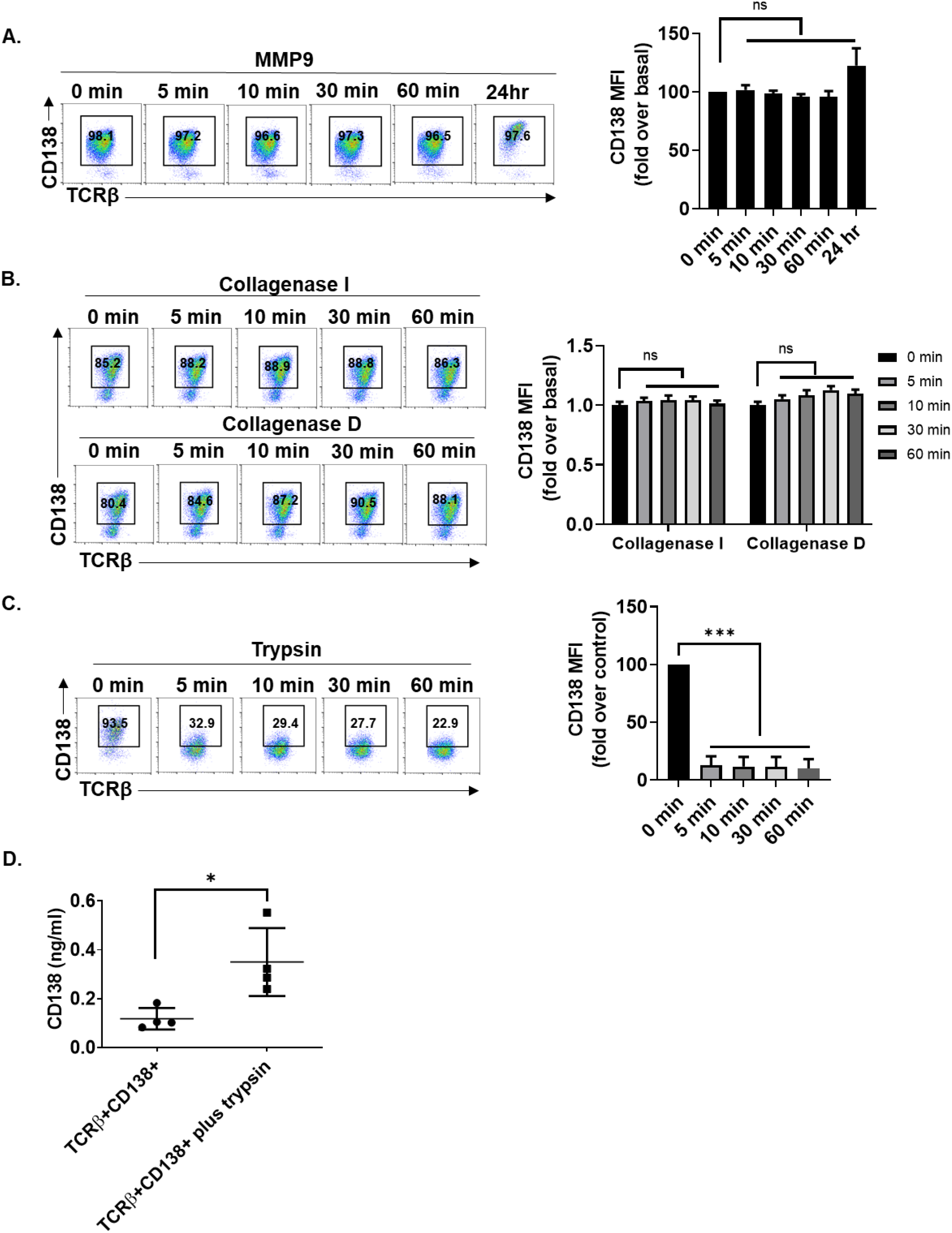
CD138, on TCRβ+CD138+ cells, is sensitive to cleavage by trypsin. Purified TCRβ+CD138+ cells were treated with active MMP9 **(*A*)**, collagenase I and D **(*B*)** or trypsin **(*C*)** for indicated duration and CD138 levels on treated cells were measured by FACS. Mean ± SD of three independent experiments are plotted. ***D***, purified TCRβ+CD138+ cells were treated with trypsin for 5 minutes, and culture supernatant CD138 levels were measured by ELISA. Mean ± SD of four independent experiments are plotted. ns, not significant, *p<0.05, ***p<0.001.

### Trypsin expressed by TCR+CD138- cells constitutively sheds CD138 from these cells

Although the pancreas is the primary source of serine protease trypsin, active trypsin is also present in mouse and human spleen cells (31). To assess whether lupus T cells also produce CD138, we measured CD138 mRNA expression in splenic T cells and found higher trypsin gene expression in purified TCRβ+CD138- cells than TCRβ+CD138+ cells (Fig. 3*A*). Also, we confirmed the higher trypsin protein production in resting TCRβ+CD138- cells by Western blot analysis (Fig. 3*B*). Moreover, αCD3/CD28 antibody (Fig. 3*C*) or PMA and ionomycin (PMA/Ion) (Fig. S3*A*, *B*) stimulation increased the levels of trypsin production in both TCRβ+CD138+ and TCRβ+CD138- cells. Coinciding with the increase in trypsin production in stimulated cells, we found a significant decrease in the expression of membrane CD138 on αCD3/CD28 antibody treated TCRβ+CD138+ cells (Fig. 3*D*). A similar decrease in CD138 expression was observed in PMA/Ion treated TCRβ+CD138+ cells (Fig. S3*C*). The simultaneous increase in the trypsin production and the decrease in CD138 expression on trypsin treated TCRβ+CD138+ cells suggest the cleavage of CD138 by the trypsin produced from TCRβ+CD138- cells. To test this hypothesis, we cultured TCRβ+CD138- cells in serum free medium and blocked the intrinsic trypsin activity by adding the trypsin inhibitors DTI or leupeptin. Although the inhibitors led to some cell death (Fig. S3*D*), TCRβ+CD138- cells expressed significantly higher CD138 after treatment with both the inhibitors (Fig. 3*E*). These results indicate that higher expression of intrinsically active trypsin in TCRβ+CD138- cells could be responsible for the shedding of CD138 from T cells in lupus mouse and contribute to the accumulation of CD138 in the MRL/Lpr mouse blood.

**Figure 3.**
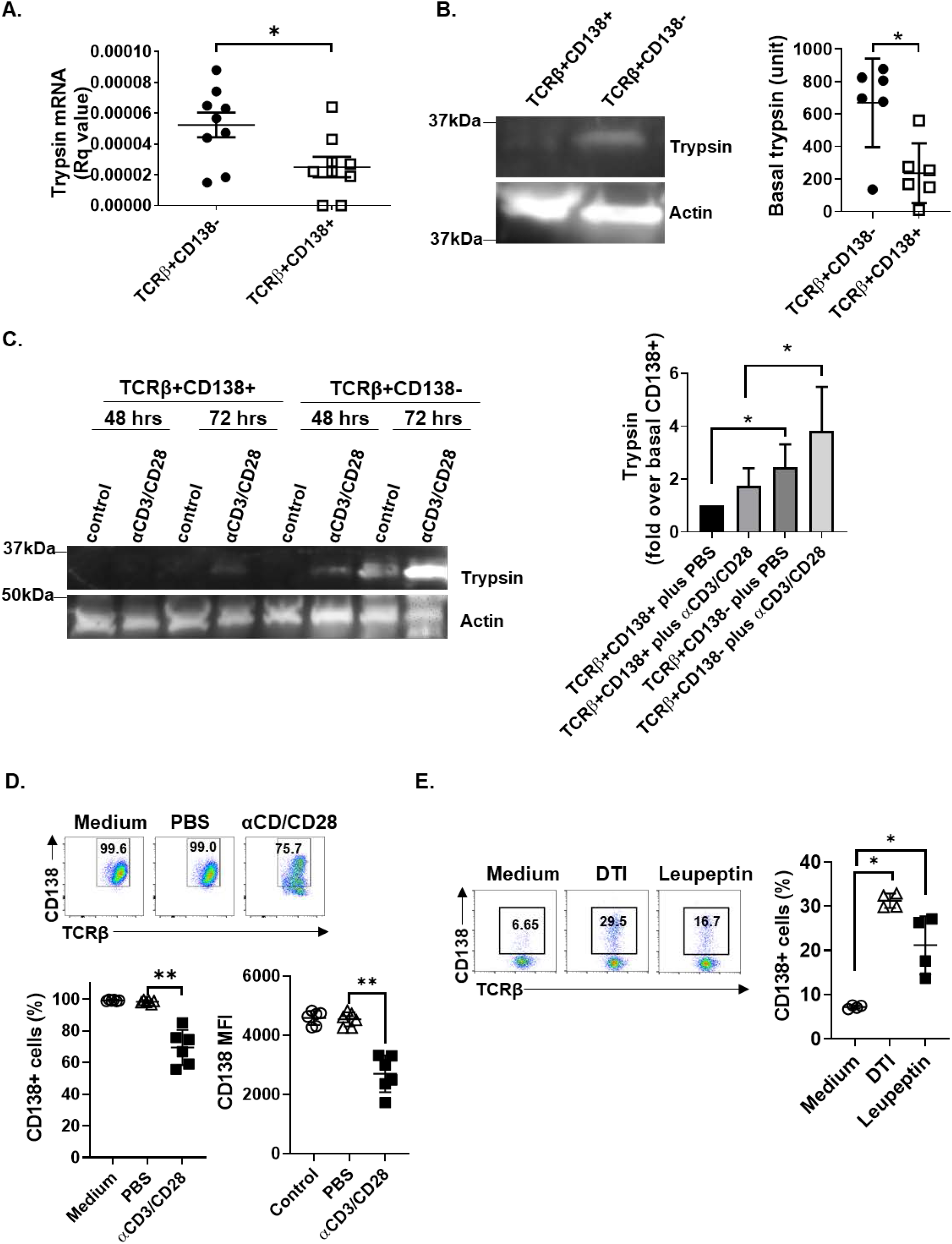
Lupus mouse splenic T cells produce biologically active trypsin. ***A***, trypsin mRNA levels were quantified in purified TCRβ+CD138+ and TCRβ+CD138- cells by Q-PCR. Mean ± SD of nine samples per group are plotted. ***B***, trypsin protein was detected in purified TCRβ+CD138+ and TCRβ+CD138- cells by Western blot analysis and the band intensities were quantified with Image J program. Mean ± SD of six different samples are plotted. ***C***, purified TCRβ+CD138+ and TCRβ+CD138- cells were treated with αCD3/CD28 antibodies for 48 or 72 hours, after which trypsin expression was detected by Western blot analysis. The band intensities were quantified with Image J program. Mean ± SD of four to five different samples are plotted. ***D***, purified TCRβ+CD138+ cells were treated with αCD/CD28 antibodies for 48 hours and membrane CD138 expression was measured by FACS. Mean ± SD of four to six different samples are plotted. ***E***, purified TCRβ+CD138- cells were cultured in serum free medium and treated with trypsin inhibitors, DTI and Leupeptin for 24 hours, after which the frequencies of CD138-expressing cells were measured by FACS. Mean ± SD of four different samples are plotted. *p<0.05, **p<0.01.

### Soluble CD138 containing serum enhances B cell differentiation and ERK activation

We next sought to investigate the function of soluble CD138 in lupus mouse serum. We first removed CD138 from serum by using anti-CD138 antibody conjugated beads (Fig. 4*A*) and tested CD138-depleted serum mediated B cell differentiation. When B cells were stimulated with APRIL and lipopolysaccharide (LPS) in CD138 depleted serum, B220^int^CD138+ plasma cell formation was significantly reduced as compared to cells cultured in serum that was treated with IgG beads only (Fig. 4*B*). The activity of CD138 on B cell differentiation was further confirmed by CD138 reconstitution experiment. Supplementation of CD138-depleted lupus mouse serum with 300 ng/ml of mouse CD138 significantly increased B220^int^CD138+ plasma cell development in APRIL and LPS stimulated B cells compared to cells stimulated with CD138-depleted serum (Fig. 4*C*). Importantly, these B220^int^CD138+ cells also exhibited elevated CXCR4 expression, an important molecule for plasma cell trafficking and maintenance (32,33). The expression of CXCR4 on plasma cells suggests that the measurement of elevated CD138 expression on B220 cells is not a result of reconstituted CD138 binding to B220 cells but is due to the differentiation of B cells into plasma cells (Fig. S4). The ERK signaling mediates key molecular switches controlling B cell differentiation (34). Since we found that lupus mouse CD138 levels increase with age and serum CD138 promotes plasma cell development, we wanted to examine whether sera from older lupus mice activate ERK pathway more than the younger mice do. Supporting this hypothesis, we found an age-dependent increase in ERK phosphorylation after treatment of B cells with sera from 3-, 8- and 14-week-old lupus mice (Fig. 4*D*). We verified the contribution of CD138 in lupus mouse serum in the ERK phosphorylation by comparing the ERK activation in B cells incubated with serum before and after CD138 depletion. Compared to unmanipulated sera, sera depleted of CD138 induced significantly less ERK phosphorylation (Fig. 4*E*). Collectively, ERK phosphorylation experiments underscore the contribution of CD138 in lupus mouse serum mediated differentiation of B cells.

**Figure 4.**
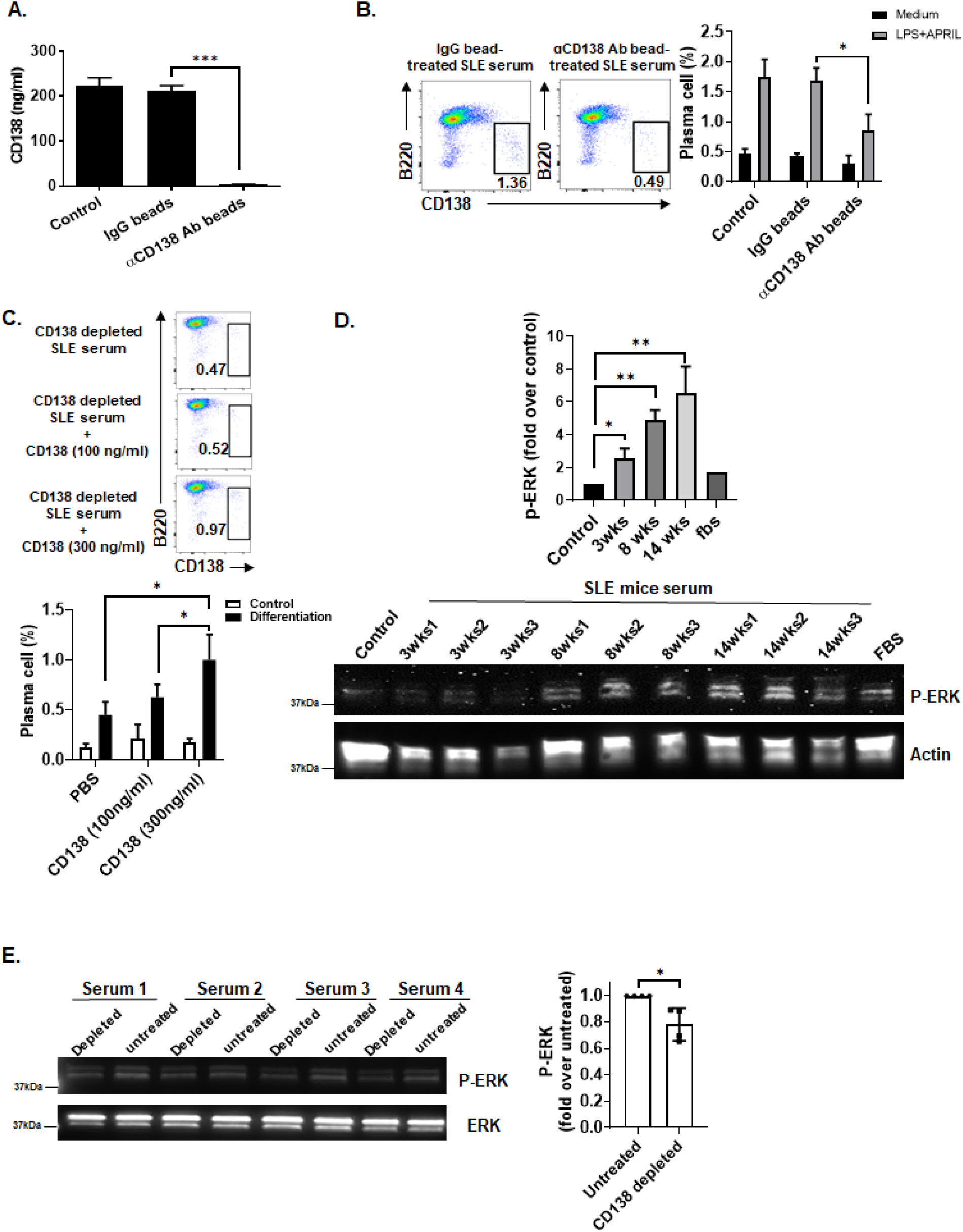
Depletion of serum CD138 in lupus mouse blunts serum induced B cell differentiation and ERK activation. ***A***, sera from 14 weeks old MRL/Lpr mice were left untreated (control), treated with Protein A/G beads coupled to IgG or CD138 antibody. The remaining soluble CD138 in the serum was quantified in ELISA. Mean ± SD of five independent experiments are plotted. ***B***, B cells from 10 to 12 weeks old lupus mice were cultured with APRIL plus LPS for 5 days in RPMI medium containing either 10% untreated (control) lupus mouse serum, lupus mouse serum pre-treated with IgG beads or anti-CD138 antibody beads. After 5 days, frequencies of plasma cells (B220^int^ CD138+) were quantified by FACS analysis. Mean ± SD of four independent experiments are plotted. ***C***, B cells from 10 to 12 weeks old lupus mice were cultured in CD138-depleted lupus mouse serum with or without addition of mouse CD138 at indicated concentrations. After 5 days of incubation, plasma cell development was assessed by FACS analysis. Mean ± SD of four independent experiments are plotted. ***D***, lupus mouse B cells cultured in RPMI medium were left untreated (control), treated with 10% FBS or with 10% serum samples from 3-, 8- and 14-weeks old lupus mice (three separate mice for each age group) for 16 hours. ERK phosphorylation was assessed by Western Blot analysis and the band intensity was quantified with Image J program. Mean ± SD of one to three independent samples are plotted. ***E***, lupus mouse B cells were incubated for 16 hours with 10% lupus mice (14 weeks old) sera that were untreated or treated with anti-CD138 antibody beads prior to incubation. ERK phosphorylation was assessed by Western Blot analysis and the band intensity was quantified with Image J program. Mean ± SD of four different samples are plotted. *p<0.05, **p<0.01.

### Soluble CD138 binds to APRIL and enhances APRIL-induced B cell differentiation and ERK activation

In CD138 transfected Jurkat T cells and CD138 positive myeloma cells, APRIL, but not BAFF, has been shown to specifically bind heparin chain of membrane bound CD138, but, to our knowledge, the binding of APRIL or BAFF to soluble CD138 has not been reported (35,36). Using CD138 antibody coated beads, we found that CD138 antibodies can pull down APRIL when CD138 and APRIL are co-incubated (Fig. S5*A*). Like serum CD138, both serum APRIL and BAFF levels increase in lupus mice as the disease progresses with age (Fig. S5*B*). Paralleling the increase in serum APRIL, we found that anti-CD138 antibody-coupled beads coimmunoprecipitated higher amounts of APRIL from the sera of older lupus mice than younger mice (Fig. 5*A*). When tested for APRIL levels, sera treated with anti-CD138 antibody coupled beads had drastically reduced levels of APRIL after the depletion of CD138 (Fig. 5*B*). Conversely and underscoring the specificity of CD138 for APRIL, serum BAFF levels remained unchanged after the pull-down with anti-CD138 antibody coupled beads (Fig. 5*B*). Thus, like its membrane bound counterpart, soluble CD138 also specifically binds to APRIL and serum CD138 is found as bound to APRIL.

**Figure 5.**
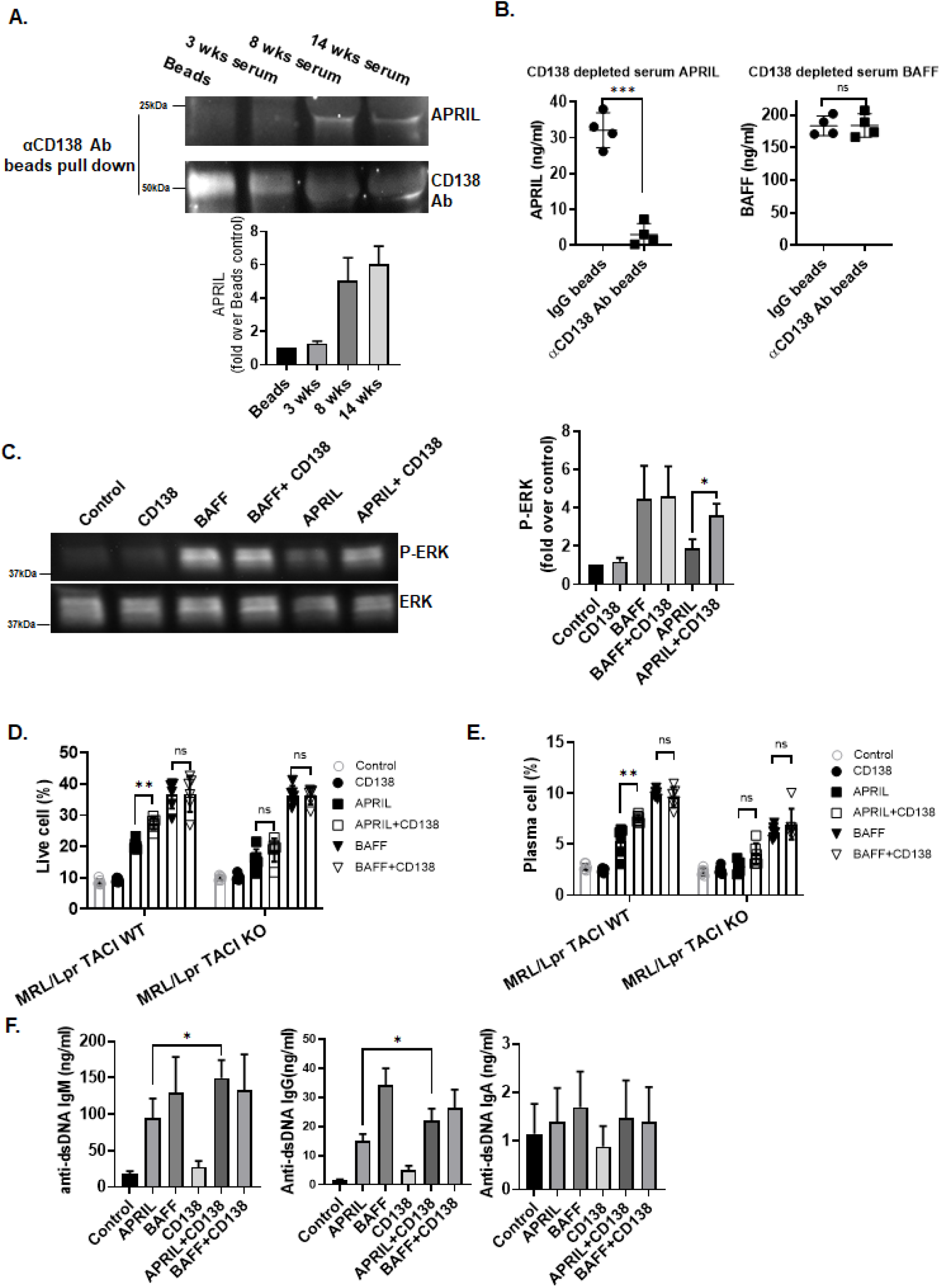
CD138 potentiates APRIL-induced ERK activation and B cell differentiation. ***A***, sera from 3-, 8- and 14-weeks old lupus mice were subjected to immunoprecipitation using CD138 specific antibody and protein A/G-Sepharose beads. A negative control precipitation without serum was run in parallel (Beads only). Precipitates were resolved by SDS-PAGE and analyzed by immunoblotting using APRIL specific antibodies as indicated. Mean ± SD of five independent experiments are plotted. ***B***, APRIL and BAFF levels were quantified by ELISA after treatment of 14 weeks old lupus mice sera in section A. Mean ± SD of 8 to 11 samples are plotted. ***C***, B cells from 10 to 12 weeks old lupus mice were cultured in 10% FBS RPMI medium for 1 hour first and then treated with APRIL, BAFF, CD138 alone or with BAFF plus CD138 and APRIL plus CD138. After overnight stimulation, ERK phosphorylation was evaluated by Western Blot analysis and the band intensities were quantified with Image J program. Mean ± SD of four independent experiments are plotted. ***D*** and ***E***, splenic B cells isolated from 10 to 12 weeks old MRL/Lpr TACI WT or MRL/Lpr TACI KO mice were cultured in 10% FBS RPMI medium for 1 hour and then treated with APRIL, BAFF, CD138 alone or in combination with APRIL and CD138 or BAFF and CD138. After 3 days, cell viability **(*D*)** and plasma cell formation **(*E*)** were assessed by FACS. Mean ± SD of six independent experiments are plotted. *F*, lupus mouse B cells were treated as in D and E. After 6 days, anti-dsDNA specific antibody levels in the culture supernatants were quantified by ELISA. Mean ± SD of four independent experiments are plotted. Ns, not significant, *p<0.05, **p<0.01, ***p<0.001.

Both APRIL and BAFF induce ERK activation (Fig. 5*C*) (37), a critical signal for B cell survival and differentiation (38,39). We next tested whether soluble CD138 modulates BAFF and APRIL induced ERK phosphorylation in B cells. Although soluble CD138 itself induced minimal ERK phosphorylation, co-stimulation of B cells with CD138 and APRIL profoundly enhanced APRIL mediated ERK activation (Fig. 5*C*). Moreover, reconstitution of CD138 depleted lupus mouse serum with CD138 led to enhanced ERK phosphorylation with APRIL (Fig. S5*C*). This potentiation effect is unique to APRIL, as combination of BAFF and soluble CD138 did not exhibit any synergy on ERK activation (Fig. 5*C*). As expected, combination of APRIL, but not BAFF, with CD138 resulted in improved lupus B cell survival (Fig. 5*D*). Both BAFF and APRIL have been shown to enhance B cell differentiation and stimulation of IgG, IgM and IgA secretion (37). Once again, while combining BAFF with CD138 did not change the percentage of plasma cells induced by BAFF, lupus B cells stimulated with APRIL and CD138 differentiated into plasma cells more than those stimulated with APRIL alone (Fig. 5*E*). Combination of APRIL with CD138 also augmented anti-dsDNA antibody production from lupus B cells (Fig. 5*F* and Fig. S5*D*). It should be noted that incubation ofB cells with CD138 and APRIL did not significantly increase total IgG, total IgA and anti-dsDNA IgA antibody production as compared to cells stimulated with APRIL alone, but it did elicit significantly higher levels of total IgM, anti-dsDNA IgM and anti-dsDNA IgG antibodies (Fig. 5*F* and Fig. S5*D*). These results indicate that binding of soluble CD138 to APRIL amplifies APRIL induced signaling, which ultimately leads to increased differentiation of autoreactive B cell into antibody secreting plasma cells in lupus mouse.

Both BAFF and APRIL share receptors TACI and BCMA on B cells (40). In primary multiple myeloma cells, membrane CD138 has been shown to function as a co-receptor for APRIL and TACI and to promote cell survival and proliferation through the APRIL/TACI pathway (35).

Consistent with these findings, although not completely, TACI deficiency significantly impaired the potentiation effect of CD138 on APRIL-stimulated B cell survival (Fig. 5*D*, Fig. S5*E*) and differentiation into plasma cells (Fig. 5*E* and Fig. S5*F*). We suspect that soluble CD138 enhances APRIL activity by oligomerizing multiple APRIL molecules. Since APRIL oligomers have higher affinity to its receptors BCMA and TACI (41), soluble CD138 bound APRIL likely induces higher ERK activation, B cell survival and differentiation through these receptors than monomeric APRIL does.

## Discussion

In this study, we reported the augmentation of APRIL mediated B cell survival, differentiation and ERK phosphorylation by soluble CD138. Our study also identified the involvement of TACI in the enhanced activity of soluble CD138 bound APRIL, as lupus B cells lacking TACI responded weaker to APRIL even when co-incubated with soluble CD138. Although, TCRβ+CD138+ cells increasingly populate lupus mouse as the disease progresses, our data strongly point to CD138 negative T cells as the primary source of soluble serum CD138 in lupus mouse because in vitro activated TCRβ+CD138- released more CD138 into the medium than TCRβ+CD138+ cells did. In support of these in vitro experiments, we have shown that lupus mice injected with TCRβ+CD138- cells accumulate more serum CD138 than those injected with TCRβ+CD138+ cells. Elevated production of soluble CD138 from TCRβ+CD138- cells was due to their high intrinsic trypsin production as membrane CD138 on lupus T cells was very sensitive to trypsin cleavage and blocking of trypsin led to CD138 retention on TCRβ+CD138- cell membrane.

Membrane CD138 is expressed at high levels in epithelial cells, plasmablasts, plasma cells and various cancer cells such as those from lung squamous cancer, adenocarcinoma, head and neck squamous cancer and mesothelioma (21,22,42,43). High level of circulating soluble CD138 has been reported in patients with multiple myeloma, lung cancer and SLE (5,6,21,44). The presence of soluble CD138 in the serum of multiple myeloma and SLE patients was believed be the result of constitutive shedding of CD138 from plasma cells (5,22). Previously, we reported the expression of CD138 on a big portion of central memory TCRβ cells in MRL/Lpr mice (23). Accumulation of these CD138 positive T cells together with the exposure of autoantigens result in the activation of host immune system and production of autoreactive antibodies in lupus mice. Despite expressing elevated levels of CD138 our study showed that TCRβ+CD138+ cells release less CD138 into the blood as compared to TCRβ+CD138- cells. In malignancies such as breast cancer and myeloma, CD138 ectodomain is believed to be shed by MMPs or collagenases produced by cancer cells (11,13,28,29). Here, we showed that these proteinases are unlikely to contribute to the cleavage of membrane CD138 from lupus T cells as CD138 on these cells was resistant to digestion by these enzymes. Instead, we found that T cell CD138 was very sensitive to trypsin treatment. Moreover, resting and activated TCRβ+CD138- cells expressed high levels trypsin, and the trypsin released from these cells effectively shed CD138 from lupus T cells in an autocrine fashion. Thus, the cleavage of CD138 from these T cell-subsets by intrinsically active trypsin is likely to be responsible for the accumulation of CD138 in the lupus mouse blood.

By binding to extracellular matrix components, integrins, growth factors, cytokines, and chemokines, membrane bound CD138 could regulate multiple biological processes, such as wound healing, cell adhesion, endocytosis, vesicular trafficking, angiogenesis and apoptosis (7). The intracellular C1 and C2 regions of membrane bound CD138 are critical for these functions as their interaction with cytoskeleton, Src kinase, or other adaptor proteins can initiate multiple downstream signaling cascades (7,42). Even though the soluble CD138, ectodomain of whole protein, lack these two vital intracellular domains, several lines of evidence indicate that soluble CD138 could also be biologically active as both competitive inhibitors and agonists. For example, soluble CD138 is shown to inhibit the mitogenicity of FGF-2, decrease the growth of carcinoma cells and induce apoptosis in myeloma cells *in vitro* (16,17,45,46). Conversely, CD138 ectodomain promotes *in vivo* tumor cell invasion as well as myeloma growth by facilitating the binding of VLA4 to VEGF receptor 2 (20,47). Similar to the function reported for membrane CD138 which provides survival advantage to myeloma or plasma cells by potentiating APRIL signaling (43), we also found an enhancement of APRIL function in inducing ERK activation and B cell differentiation when soluble CD138 is present. Importantly, soluble CD138 did not provide a detectable enhancement of BAFF activity as no increase in cell survival, differentiation and antibody secretion was observed in B cells treated with BAFF in the presence of CD138. Moreover, as in CD138 expressed on membrane of HEK293 cells, Jurkat cells and primary multiple myeloma cells, which exhibit strong binding to APRIL but not to BAFF (35,36), we also observed co-precipitation of APRIL with soluble CD138. Although we do not know the exact number of APRIL molecules binding to CD138, previous reports indicated that thousands of epidermal growth factors and APRIL molecules can bind to membrane syndecan-1 through its heparan sulfate chain (35,48). Thus, the enhancement of APRIL activity by soluble CD138 is likely a result of the oligomerization of CD138 bound APRIL molecules.

Previous studies reported the binding of membrane CD138, syndecan-2 and syndecan-4 to TACI (35,49). It is possible that soluble CD138 can augment APRIL activity by simultaneously binding to both APRIL and its receptor TACI. Suggesting a partial involvement by TACI, the enhancement mediated by CD138 on APRIL induced signaling, cell survival and autoreactive antibody production was decreased but not totally abolished in TACI deficient MRL/Lpr mice. Since APRIL engages BCMA in addition to TACI, and BCMA is involved in plasma cell generation and maintenance (50,51), both TACI and BCMA likely mediate APRIL oligomer activity.

Both SLE patients and lupus mouse present with an increase in blood CD138 as the disease progresses. Building on our recent report where we studied the involvement of TCRβ+CD138+ cells in lupus immunopathogenesis (23), we now show that CD138 released from TCRβ+CD138- cells contribute to the increase in circulating CD138 pool. Our study also suggests a plausible role for soluble CD138 in augmenting APRIL induced activation of autoreactive B cells in lupus. These findings highlight a previously unappreciated participation of CD138 in lupus pathogenesis, which may be exploited to evaluate disease progression and/or may unveil novel therapeutic targets.

## Experimental Procedures

### Mice

MRL/MpJ-FASLPR/J (referred to as MRL/Lpr throughout the manuscript), were purchased from The Jackson Laboratory (Bar Harbor, ME). TACI deficient MRL/Lpr mice (MRL/Lpr TACI KO) and the wild-type counterpart (MRL/Lpr TACI WT) were described previously (26). All mice were bred and housed under specific pathogen-free conditions in the animal facility of US Food and Drug Administration (FDA)/Center for Biologics Evaluation and Research (CBER) Veterinary Services. The use of animals was approved by and carried out within accordance of the US FDA/CBER Institutional Animal Care and Use Committee (permit numbers 2002–31 and 2017-47). All methods were performed in accordance with the relevant guidelines and regulations.

### Flow cytometry

Single cell suspensions of spleens were prepared by mechanical dissociation of tissue through a 40 μM cell strainer. Red blood cells were then lysed using ACK lyses buffer (Lonza, Wallersville, MD). Mouse blood leukocytes were collected by lysing red blood cells with ACK buffer and centrifugation at 300 x g for 5 minutes. Cells were stained with fluorescent labelled anti-mouse antibodies after blocking CD16/CD32 with Fc Block (BD Biosciences, San Jose, CA). Flow cytometry analysis was performed using following antibodies: Pacific blue anti-CD19, BV421 anti-CD19, APC anti-TCRβ, APC CXCR4, Percp Cy5.5 B220 (All from BioLegend, San Diego, CA), and PE–anti-CD138 (from BD biosciences). DAPI and LIVE/DEAD™ Fixable Near-IR Dead Cell Stain Kit were purchased from Thermo Fisher Scientific (Waltham, MA). APRIL-Alexa488 and BSA-Alexa488 were prepared by conjugation of APRIL and BSA to Alexa Fluor 488 NHS ester (Thermo Fisher). Stained cells or beads were analyzed using a flow cytometer (LSR II; BD) and data were analyzed using FLOWJO, version 10.1 for PC (Tree Star, Ashland, OR).

### Measurement of CD138, APRIL, BAFF production

CD138 levels in MRL/Lpr mice serum or culture supernatants were measured using mouse SDC1 ELISA Kit from Aviva Systems Biology (San Diego, CA) according to the manufacturer’s instructions. Serum APRIL and BAFF levels in MRL/Lpr mice blood were measured using mouse APRIL ELISA kit (Mybiosource, San Diego, CA) and mouse BAFF/BLyS/ TNF13B DuoSet ELISA kit (R&D Systems), respectively.

### MMP9 activity measurement

Recombinant mouse MMP9 protein was purchased from Abcam (Cambridge, MA). To obtain maximum latent MMP9, MMP9 was treated with 2.5 mM 4-Aminophenylmercuric acetate (APMA, Sigma-Aldrich) in a buffer containing 50 mM Tris-HCL pH 7.5, 1 mM CaCl_2_, 0.05% Triton X-100 at 37°C for 1 hour prior to use. The activity of MMP9 was verified with MMP9 colorimetric drug discovery kit (Enzo, Farmingdale, NY) according to the manufacturer’s instructions.

### T cell isolation and stimulation

MRL/Lpr mice splenic T cells were purified with Dynabeads™ FlowComp™ Mouse Pan T (CD90.2) Kit and dissociated from beads according to manufacturer’s instructions (ThermoFisher). After staining purified T cells with PE anti-CD138 antibody, TCRβ+CD138- and TCR+CD138+ cells were further separated with anti-PE magnetic microbeads (Miltenyi Biotec, Auburn, CA). Depending on the experimental objective, purified TCR+CD138+ and TCR+CD138- cells were stimulated with 1 μg/ml anti-CD3 and -CD28 antibodies (BD Pharmingen), 10 ng/ml PMA and 100 ng/ml ionomycin, 100 mg/ml Collagenase I, 100 mg/ml Collagenase D, 2.5 mg/ml trypsin, 10 μM leupeptin (all from Sigma-Aldrich, St. Louis, MO), 10 μg/ml MMP9, TrypLE or Defined Trypsin Inhibitor (DTI) (all from ThermoFisher) for indicated duration, and the expression levels of cell surface CD138 were quantified by FACS. Trypsin gene expression levels were quantified by Q-PCR and trypsin protein was analyzed by Western Blot using rabbit polyclonal trypsin antibody (ThermoFisher).

### T cell adoptive transfer experiments

Purified TCRβ+CD138- and TCRβ+CD138+ cells were washed three times with PBS before resuspending them in PBS. MRL/Lpr mice were injected i.v. with 1 x 10^7^ cells in 100 μl of PBS.

### B cell isolation and stimulation

Splenic B cells were isolated from 10 to 12 weeks old MRL/Lpr mice using B Cell Isolation Kit (Miltenyi Biotec). Isolated cells were washed three times with PBS, after which their purity was assessed by flow cytometry (purity was greater than 97%). For B cell stimulation and survival, 1 x 10^6^ B cells were cultured in RPMI media containing 10% FBS or 10% of anti-CD138 beads pretreated or IgG coated beads pretreated MRL/Lpr mice serum. Next, cells were stimulated with 500 ng/ml of recombinant APRIL (Peprotech, Rocky Hill, NJ) or recombinant BAFF (R&D Systems) with or without recombinant mouse CD138 (R&D Systems) for 24 or 48 hours. Phosphorylation of ERK was assessed in Western Blot analysis and the survival of cells was analyzed by FACS after DAPI staining. For B cell differentiation analysis, B cells were first cultured in RPMI media containing 10% anti-CD138 beads pretreated MRL/Lpr mice serum with, and then were incubated with 250 ng/ml of APRIL and 10 ng/ml of LPS for 5 days. Stimulated cells were harvested, and percentages of plasma cells were assayed on DAPI negative cells with FACS after staining with B220, CD138 and CXCR4 antibodies. Culture supernatants were also collected and total or anti-dsDNA specific IgM, IgG and IgA antibody concentrations were determined in ELISA as described previously (26).

### Quantitative Real-Time PCR

Total RNA was extracted from the FACS sorted cells using RNeasy Mini kit (Qiagen, Germantown, MD). Two hundred nanograms of total RNA were reverse transcribed into cDNA using random hexamers with the Taqman Reverse transcription kit (Invitrogen). The expression of targeted genes and GAPDH were determined using Taqman Gene Expression assays and CFX96 Touch Real-Time System (BioRad, Hercules, CA). Relative expression values were determined by the 2-ΔCt method where samples were normalized to GAPDH expression as described previously (52).

### Statistical analysis

Data from groups were compared using GraphPad Prism, Version 8 software (GraphPad Software, San Diego, CA) and nonparametric testing was performed by the Mann-Whitney rank sum test for two groups and by Kruskal-Wallis two-way ANOVA on ranks for three or more groups.

## Supporting information

This article contains supporting information.

